# SiCR: Web application for single-cell repertoire analysis and immune profiling

**DOI:** 10.1101/2023.08.03.551897

**Authors:** Masakazu Ishikawa, Kaoru Matsumoto, Daisuke Okuzaki

**Author notes:** Corresponding author: Daisuke Okuzaki.

## Abstract

**Background:** Single-cell RNA sequencing (scRNA-seq) allows analysis of complete sequences of antigen receptors in individual cells. However, it is a complex technique that requires multiple analyses to obtain accurate results. Although several user-friendly tools for scRNA-seq are available, none are specifically designed for immune profiling.

**Results:** We developed a web application called SiCR that is based on the Shiny framework of the R package and specializes in single-cell immune profiling. SiCR allows clustering and cell typing required for both general single-cell and immune profiling analyses, such as predicting whether the chronotype is expanding in each group and the antigen the expanding chronotype targets. These analyses can be performed using a cursor control. SiCR also allows for detailed figure settings, enabling immediate publication of results.

**Conclusions:** SiCR is a comprehensive workbench that can be used by biologists for single-cell immune profiling. Currently, it is the only web application that allows single-cell repertoire analysis using both raw and preprocessed data. Moreover, SiCR significantly reduces the time and effort required to analyze and interpret information in single-cell immune profiling. Therefore, SiCR is a potential reference application for interactive analysis and investigation of biological data, especially for immune profiling.

## Background

The diversity of T (TCR) and B (BCR) cell receptor gene repertoires constitute the core of the adaptive immune system and serves as an essential defense mechanism against invasion by viruses, bacteria, and other pathogens. TCRs are composed of TCR alpha (TRA) and beta (TRB) chains, whereas BCRs contain immunoglobulin light/kappa (IGL/K) and IG heavy (IGH) chains [1]. The diversity of TCRs and BCRs is vast because of the process of V(D)J gene rearrangement and random insertion and deletion of non-target nucleotides of uncertain lengths between V-D and D-J (TRB, IGH) or V-J junctions (TRA, IGL/K) [2]. High-throughput sequencing using various platforms, including Roche 454 and Illumina HiSeq, has been applied to repertoire analysis [3–7].

For example, high-throughput sequencing of BCRs has shown promise in elucidating the human adaptive immune response, including the sequence of mutations that occur during coevolution of antibodies with human immunodeficiency virus [8]. Moreover, the development of high-throughput sequencing technologies such as single-cell RNA sequencing (scRNA-seq) has allowed us to analyze nucleic acids of individual cells [9]. Single-cell RNA-seq enables acquisition of full-length variable regions of both H and L chains from individual T/B cells, while maintaining a pair of chain information at scale [10–12]. Single-cell RNA-seq has led to the development of several computational tools to address various aspects of data analysis [13]. However, dependency on computational knowledge poses a significant barrier that affects researcher ability to explore data, especially for those with limited expertise in command-line interfaces. However, multiple analyses are necessary because the results of each step in the analysis is significantly influenced by data quality and execution parameters. To re-evaluate the data, various tools must be easily accessible to biologists. Several web application tools have recently been developed for scRNA-seq [14–18]. These tools aim to solve similar problems by providing graphical user interfaces instead of command-line interfaces, often addressing various aspects of analysis, including diverse content. However, none of these tools cover single-cell repertoire analysis.

This paper describes the interactive web application SiCR that we developed for single-cell repertoire analysis. SiCR provides a comprehensive single-cell repertoire analysis workbench. The architecture and capabilities of SiCR are described based on analysis of a publicly available dataset.

## Implementation

### Overview

SiCR uses the R programming language and Shiny framework to create an easy-to-use interface. SiCR has different parts that perform various analyses, including: (1) normalizing and clustering using the Seurat software [19]; (2) combining information on TCRs/BCRs from each cell using the scRepertoire package [20]; (3) calculating gene usage by TCRs/BCRs using the tidyverse package [21] and Alakazam [22]; (4) analyzing the diversity of TCRs/BCRs, such as alpha diversity [23-25] and clonotype expansion; (5) predicting antigens that TCRs/BCRs target, using existing databases; and (6) creating a phylogenetic tree of the same clonotypes (BCR only) using Dowser [26].

### Input

SiCR requires a maximum of three input files generated using the Cell Ranger software of 10x Genomics. When TCR/BCR analysis was not required, the cellranger count pipeline was used for the analysis, whereas the cellranger multi pipeline was used if TCR and/or BCR data were available. If multiple runs needed to be merged into a single file, the cellranger aggr pipeline was used. While analyzing data using the cellranger multi pipeline, the metadata, such as donor information, can be input as a comma-separated value (CSV) file, which can be used in the analysis using SiCR. The required input files for SiCR consist of a mandatory HDF5 file, which is the sample_filtered_feature_bc_matrix.h5 file generated by Cell Ranger, and two optional TCR or BCR-filtered _contig_annotations.csv files.

### Clustering and visualization

The clustering module of SiCR is based on the Seurat package (v4) [19] and provides a rich graphical user interface using the Shiny framework of the R package, enabling interactive data manipulation and execution of Seurat functions. When output files of Cell Ranger (10x Genomics) are uploaded, SiCR automatically performs the following steps before displaying the results: data normalization, dimensionality reduction through principal component analysis, clustering of cell populations, and annotation. The clustering results are visualized as a uniform manifold approximation and projection (UMAP) and bar plot for each fraction. The UMAP and bar plot can be grouped by the Seurat cluster, sample number, annotation, and metadata set using the cellranger aggr pipeline. Moreover, the sizes, labels, and legends of the graphs and dots can be customized. Additionally, users can download the results as high-resolution plots or CSV files. Automatic annotation is performed using ScType [27], which determines the positive and negative genes for each subset, assigns scores to every single cell, and further assigns the most highly scored subtype to each cell among the same seurat_cluster.

### Gene usage

SiCR allows its users to visualize the gene usage of each cell group, providing insights into the biases and preferences of the immune repertoire. Users can compare the gene usage between the cell groups in absolute or relative amounts. Specifically, for TCR analysis, users can compare the usage of the TRA V and J genes, as well as the TRB V, D, J, and C genes. For BCR analysis, users can compare the usage of IGH V, D, J, and C genes as well as that of IGL/K V and J genes. CSV files are generated for all comparisons.

### Clonotype expansion

Clonotype expansion is one of the most interesting areas of study in repertoire analysis. SiCR can help visualize the expansion of TCR and BCR clonotypes for each group, as defined by Cell Ranger. The number of clones is displayed as a bar plot or heatmap in absolute or relative amounts (100%). Additionally, SiCR presents the abundance and diversity curves that are provided by Alakazam [22]. SiCR also uses alpha diversity, as proposed by Hill [23], to calculate and display a smooth curve. The diversity index for each group was the average value over all resampling realizations, and confidence intervals were derived using the standard deviation of the resampling realization, as described by Chao et al. [25]. The significance of the difference in the diversity index between groups was tested by constructing a bootstrap delta distribution for each unique pair of values in the group column.

### Antigen prediction

Antigen prediction is crucial in repertoire analysis, as it allows for a deeper understanding of how the immune system reacts to various antigens. SiCR uses VDJDB [28], McPAS-TCR [29], and PIRD [30] for the TCR database and PIRD and Scepter [31] for the BCR database. These databases provide the sequences of the CDR3 region of TRB or IGH, as well as information on the antigens that interact with the CDR3 region. When the amino acid sequence of CDR3 of TRB or IGH in the single-cell sequencing data completely matches the amino acid sequence of CDR3 from the CDR dataset listed in the database, information on the antigen that interacts with CDR3 is returned.

### Phylogenetic analysis (only BCR)

Upon recognition of an antigen, B cells undergo proliferation and hypermutation of BCRs [32]. Somatic hypermutation increases BCR diversity. Phylogenetic analysis of BCR sequence evolution over time can be tracked, and the development of immune responses and immunological memory can be studied by displaying cells with the same clonotype in a phylogenetic tree. Cell Ranger (versions 5.0 or higher) groups B cells into clonotypes using a module for clonal analysis called enclone [33] that filters and groups cells into clonotypes simultaneously. SiCR uses a tool called Dowser [26,34] to display the phylogenetic trees and clonotypes identified by enclone. Users can select the clonotype number and create a phylogenetic tree of cells identified with the same clonotype to track B cell mutation. Only the clonotypes with two or more contigs were selected. Once a clonotype is selected, a phylogenetic tree is created using contigs recognized by the same clonotype, and the VH region of the germline is created with reference to the contig. The phylogenetic tree is constructed using the maximum likelihood method [35].

## Results

To demonstrate the functionality of SiCR, we analyzed the peripheral blood mononuclear cell (PBMC) data from Sureshchandra et al. [36] publicly available in 10x Genomics. Sureshchandra et al. conducted scRNA-seq of enriched T/B cells after two doses of mRNA vaccination, with responses observed in convalescent individuals with asymptomatic disease. Here, after running Cell Ranger, we obtained 50,147 cells. Among them, 34,920 cells contained TCR information and 9,399 contained BCR information. Figure 1A shows the UMAP of the samples analyzed using SiCR. For our analysis, only T and B cells were sorted. Therefore, SiCR automatically annotated most cells as T or B cells. Figure 1B shows the proportion of subsets for each condition. As shown in Figure 1B, the proportion of naïve B cells was higher in HCW_baseline than in HCW_convalescent patients. The TCR analysis data is presented in Fig. 2. First, based on the results of alpha diversity analysis, diversity decreased as follows: HCW_convalescence > Vaccine_post > Vaccine_pre > HCW_baseline (Fig. 2A). This finding is consistent with the results of the previous study [36]. The clonal abundance curve showed that HCW_convalescence had a steep slope, indicating the expansion of a specific clonotype (Fig. 2B). The heatmap showed that in the HCW_convalescence group, clonotype 1 accounted for 10% of all TCRs and was the most expanded (Fig. 2C). Clonotype 1 accounted for 2.7% of all TCRs in the HCW_baseline group, although the most abundant clonotype was clonotype 3, accounting for 4.2% of all TCRs. Antigen prediction was performed based on CDR3 of TRB for these two clonotypes. No corresponding antigen was detected for clonotype 1, but clonotype 3 exhibited reactivity with cytomegalovirus IE1 (Fig. 2D). Figure 3 presents the BCR data. Regarding alpha diversity under the condition of q = 1, also called Shannon diversity, the HCW_convalescent group had a higher diversity score than the HCW_baseline group, and the Vaccine_pre group had a higher diversity score than the Vaccine_post group (Fig. 3A). Regarding clonotype abundance, expanding clonotypes were observed in the HCW_baseline group, but the degree of expansion was lower than that in TCRs (Fig. 3B). In addition, regarding the proportion of C gene usage by IGH, the proportion of IGHM (immunoglobulin heavy constant mu) was lower in the HCW_convalescent and Vaccine_pre groups than in the HCW_baseline and Vaccine_post groups, whereas that of IGHG1 (immunoglobulin heavy constant gamma 1)was higher in the HCW_convalescent group and those of IGHA1 and IGHA2 (immunoglobulin heavy constant alpha 1 and 2) were higher in the Vaccine_pre group (Fig. 3C). Based on the investigation to ascertain the most expanded clonotype in the HCW_baseline group, clonotype 1 accounted for 0.5% of all BCRs (Fig. 3D). This clonotype was divided into two sub-clonotypes, and the differences from the germline were similar (Fig. 3E).

**Figure 1.**
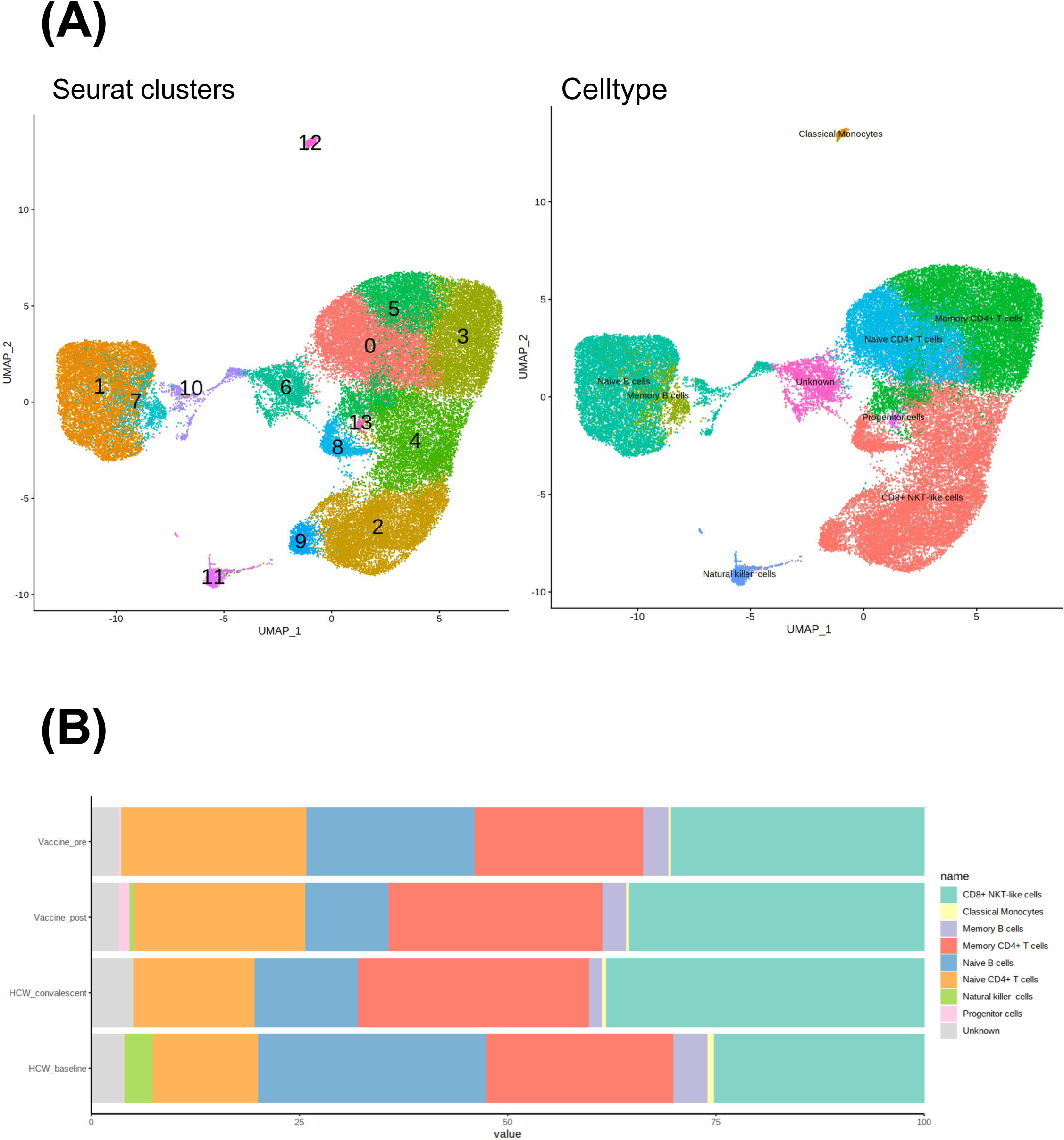
Figures displayed in the “Gene expression” section of SiCR. **a)** UMAP dimensionality reduction embeddings of PBMCs, colored based on the Seurat clusters, origin, and cell types. **b)** Cell type proportion for each origin.

**Figure 2.**
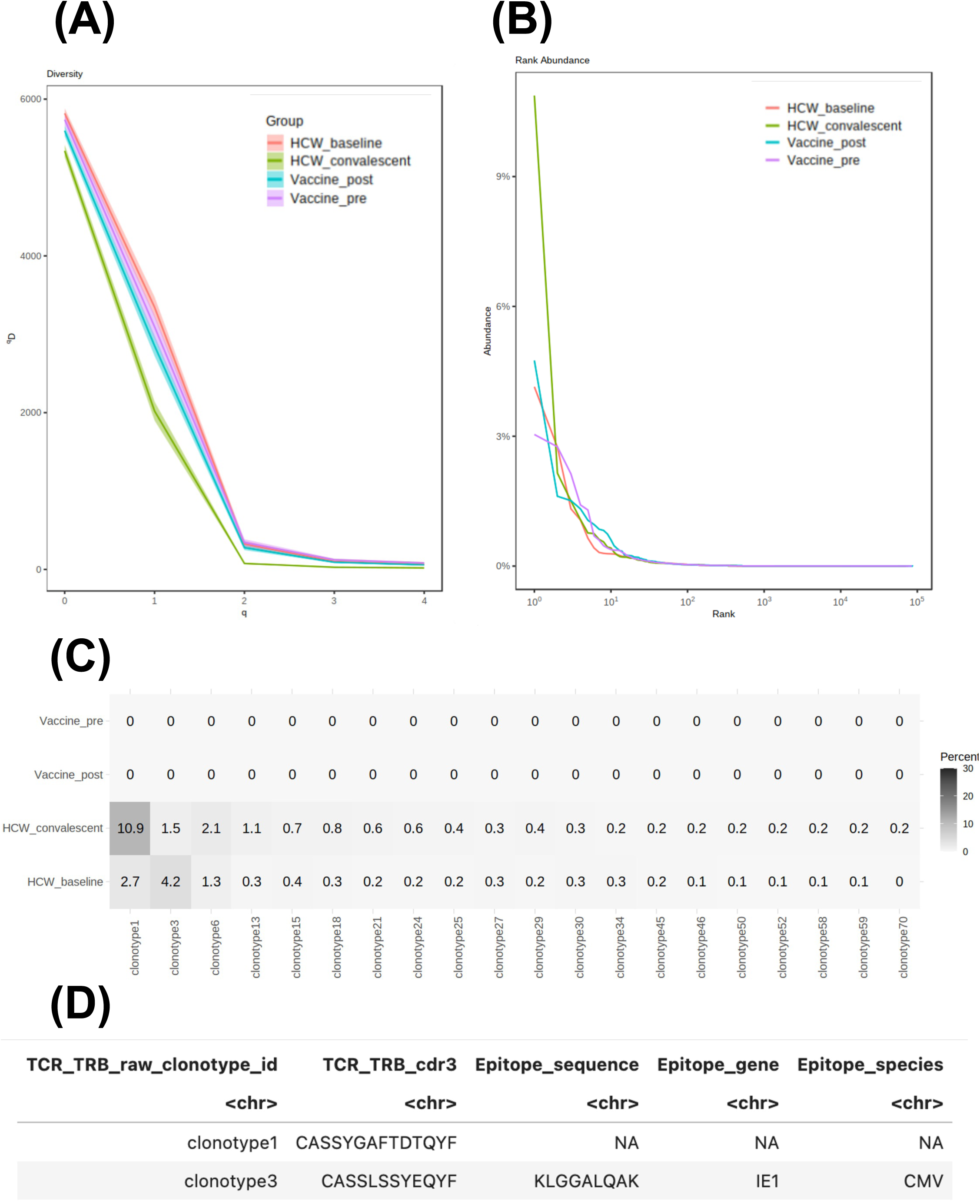
Figures displayed in the “TCR” section of SiCR. **a)** Alpha diversity curve of the TCR clonotypes, grouped by their origin. **b)** Clonal abundance curve of the TCR clonotypes, grouped by their origin. **c)** Heatmap describing the expanded clonotypes arranged in order of the most to least abundant clonotype in the HCW_convalescent group. **d)** The predicted antigen interacts with the TRB CDR3 region of either clonotype 1 or clonotype 3.

**Figure 3.**
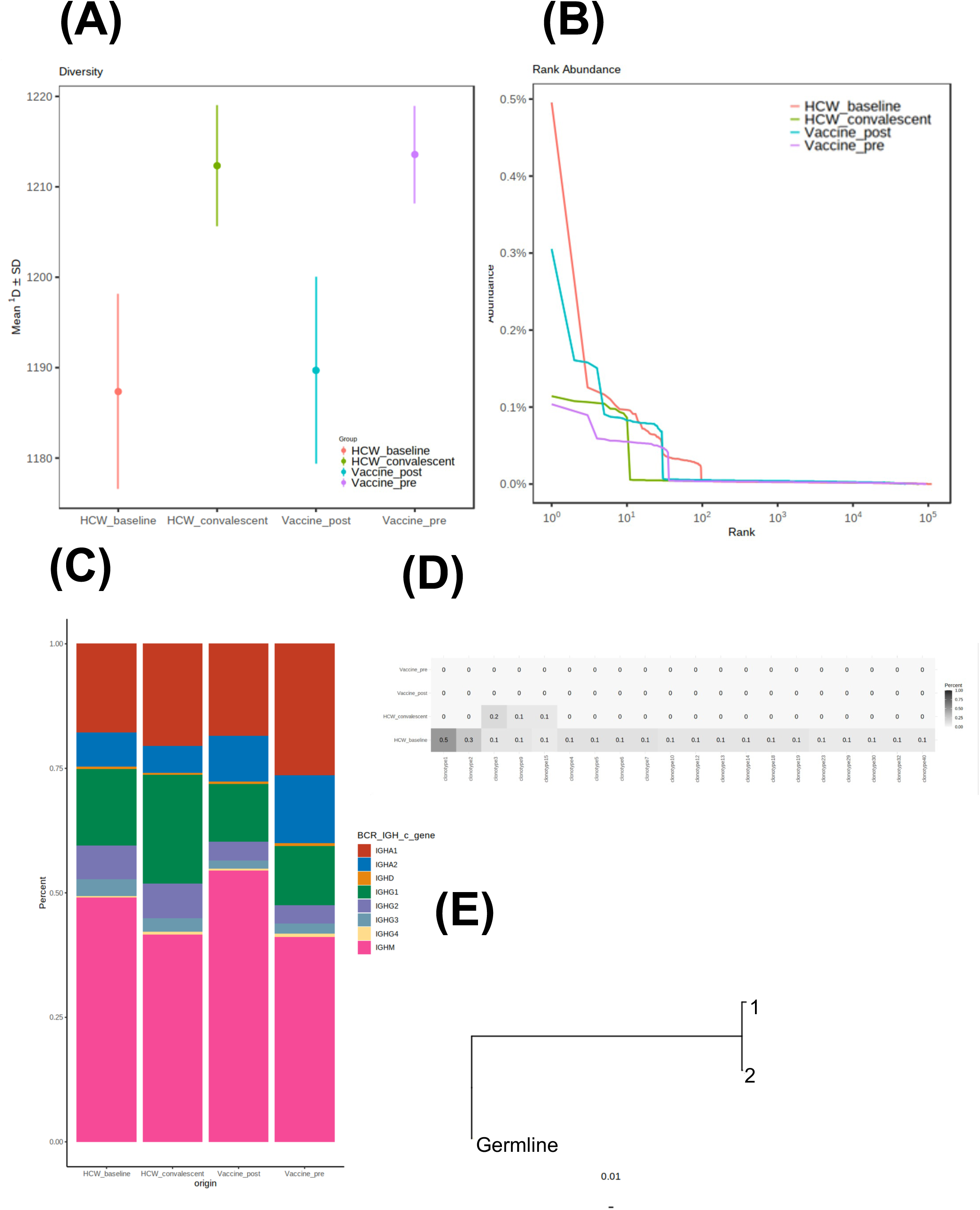
Figures displayed in the “BCR” section of SiCR. **a)** Alpha diversity of the BCR clonotypes, grouped by their origin. **b)** Clonal abundance curve of the BCR clonotypes, grouped by their origin. **c)** Proportion of the C gene of IGH in each origin. **d)** Heatmap describing the expanded clonotypes arranged in order of the most to least abundant clonotype in the HCW_baseline group. **e)** Phylogenetic tree of clonotype 1 rooted by the germline.

## Discussion

SiCR is a bioinformatics pipeline that can perform multiple previously reported single-cell repertoire analyses. SiCR assists researchers in deciphering complex biological mechanisms conveniently through cutting-edge analysis modules and visualization functions. To provide a basis for comparison, we compiled a catalog of similar tools (SCALA [37], Asc-Seurat [14], Bingle-seq [18], SingleCAnalyzer [38], Cerebro [39], Is-CellR [40], and WASP [41]) and highlighted their strengths and weaknesses along with their complementarity to SiCR (Table 1). To the best of our knowledge, SiCR is the only web application that allows single-cell repertoire analysis. Therefore, we expect SiCR to become a reference application to interactively analyze and explore data, owing to its simplicity and usefulness.

Although SiCR can accelerate single-cell repertoire analysis, it has certain limitations. First, its application is currently limited to human samples. This is because its automatic annotation targets data from human samples. However, the immune profiling kit of 10x Genomics can now handle mouse samples in addition to human samples; therefore, SiCR should be able to handle data from mouse samples soon. Second, the automatic annotation of the samples is scored based on whether each gene is positive or negative. Therefore, although accurate annotation can be performed for diverse cells such as PBMCs, there is a possibility of error when diversity is reduced, such as when only T or B cells are used. Third, there is a memory-related problem: SiCR can be run in a localized system but requires a large amount of memory for clustering. To address this, we plan to implement SiCR on a public server. In addition, scRNA-seq analyses other than repertoire analyses, such as differential gene expression and trajectory analyses, have not been implemented in SiCR yet. These analyses can be performed using the Loupe browser provided by 10x Genomics or other web tools; therefore, they were not implemented in SiCR. However, it would be ideal if all the analyses could be performed using SiCR in the future.

## Conclusion

SiCR is a powerful bioinformatics pipeline for single-cell repertoire analysis, offering a range of cutting-edge analysis modules and visualization functions. It is capable of handling both raw and preprocessed data, allowing users to perform analyses from scratch or reanalyze preprocessed data. Moreover, SiCR is the only web application that allows single-cell repertoire analysis and has the potential to become a reference application to interactively analyze and explore data owing to its simple interface and utility.

## List of abbreviations

TCR: T cell receptor
BCR: B cell receptor
TRA: TCR alpha
TRB: TCR beta
IGL/K: Immunoglobulin light/kappa
IGH: Immunoglobulin heavy
scRNA-seq: Single-cell RNA sequencing
CSV: Comma-separated value
PBMC: Peripheral blood mononuclear cell

## Availability and requirements

**Project name:** SiCR (Web application for single-cell repertoire analysis)

**Project home page:** https://github.com/Masakazu-Ishikawa137/SiCR

**Operating system(s):** Platform independent

**Programming language:** R (Shiny)

**Other requirements:** R packages ggsci, RColorBrewer, tidyverse, Seurat, shiny, HGNChelper, alakazam, dowser, hdf5r, BiocManager, openxlsx, hrbrhemes, Biostrings, GenomicAlignments, ggtree.

**License:** GPL - 3.0

**Any restrictions to use by non-academics:** None.

## Declarations

### Ethics approval and consent to participate

Not applicable.

### Consent for publication

Not applicable.

### Availability of data and materials

The dataset analyzed in the study was generated by Sureshchandra et al. [36] on NCBI’s Sequence Read Archive under BioProject PRJNA767017.

### Competing interests

The authors declare that they have no competing interests.

### Funding

This work was supported by the Japan Agency for Medical Research and Development (AMED) (grant number 20fk0108404h0001), the Japan Agency for Medical Research and Development–Core Research for Evolutional Science and Technology (AMED– CREST) (22gm1810003h0001), the Mitsubishi Foundation, and the NIPPON Foundation for social innovation.

### Authors’ contributions

M.I. participated in the design, testing, and implementation of the web application, design and implementation of the web page containing SiCR’s documentation, and in the writing of the manuscript. K.M. participated in the web application implementation. D.O. tested the web application in the multiple stages of its implementation and participated in the web application design, supervision of its implementation, and the manuscript’s writing. All authors read and approved the final manuscript.

## Acknowledgements.

Not applicable.

## Authors’ information

**(optional)**

## Notes

### Competing Interest Statement

The authors have declared no competing interest.

